# Multi-Scale Machine Learning for Antibody-Antigen Binding Affinity Prediction Using Deep Mutational Scanning and Structural Features

**DOI:** 10.64898/2026.06.09.730151

**Authors:** Santhosh Sivasubramani

## Abstract

Predicting how mutations alter antibody-antigen binding affinity is essential for antibody engineering and vaccine design, yet current methods generalize poorly to unseen complexes. We present a multi-scale machine learning framework integrating 93 descriptors across four modalities: physicochemical, structural, ESM-2 protein language model, and solvent-accessible surface area (SASA)/ΔΔ*G*_fold_ features. Under leave-one-complex-out deep mutational scanning (LOCO-DMS) cross-validation on AbAgym (36,541 mutations, 68 experiments, 13 pathogens), gradient boosting achieved MCC = 0.206; a confidence-stratified ensemble reached MCC = 0.374 (83.5% accuracy, 25.5% coverage). No single modality exceeds the majority baseline alone; only multi-scale fusion succeeds. Boltzmann ceiling analysis shows 45.9% of mutations are near-neutral (|ΔΔ*G*| *< k_B_T*), bounding theoretical maximum MCC at 0.473; our method achieves 79.1% of this limit. Five deep learning architectures benchmarked under LOCO-DMS showed self-attention matching gradient boosting (MCC = 0.200). Cross-pathogen transfer failed systematically (mean 46.7%), confirming universal binding predictors remain an open challenge.

## Introduction

Antibody-antigen interactions, mediated by the binding of antibody complementarity-determining regions (CDRs) to specific epitopes on an antigen surface, are central to adaptive immunity and underpin the development of therapeutic antibodies for cancer, infectious diseases, and autoimmune disorders [1, 2]. The ability to predict how mutations affect binding affinity is essential for antibody engineering, understanding viral immune escape, and designing broadly neutralizing vaccines [3, 4]. Despite substantial progress in computational methods, accurate prediction of binding affinity changes (ΔΔ*G*) upon mutation remains an open challenge, particularly for mutations with subtle effects on binding [5].

The Structural Kinetic and Energetic database of Mutant Protein Interactions (SKEMPI) has served as the primary benchmark for binding affinity prediction methods, containing thermodynamic measurements for over 7,000 mutations across 345 protein complexes [6]. Methods evaluated on SKEMPI report strong correlations between predicted and experimental values; however, concerns have emerged regarding data leakage in conventional cross-validation schemes, where training and test sets share mutations from the same protein complex [5]. Leave-one-complex-out (LOCO) cross-validation provides a more stringent evaluation of generalization to unseen protein systems.

Recent advances in deep learning have enabled new approaches to binding affinity prediction. Protein language models trained on millions of sequences, such as Evolutionary Scale Modeling 2 (ESM-2) [7], capture evolutionary constraints that inform functional prediction. Graph neural networks (GNNs) represent protein structures as graphs and learn representations that capture local geometric features [8, 9]. Deep mutational scanning (DMS) experiments provide comprehensive mutational landscapes for specific antibody-antigen systems [10], with databases such as AbAgym aggregating over 300,000 mutations across diverse pathogens [11].

Despite these advances, fundamental questions remain regarding the limits of predictability for binding affinity changes. Near-neutral mutations, where thermal noise approaches the magnitude of the binding energy change, may be intrinsically unpredictable from sequence or structure alone [12, 13]. Understanding these limits is essential for appropriate application of computational predictions in drug discovery. Recent work has highlighted this challenge: Hummer et al. [14] showed that orders of magnitude more data may be needed for generalizable ΔΔ*G* prediction, while Janusz et al. [15] demonstrated “poor generalization of binding predictions from machine learning models” on systematically generated mutant data. Critically, existing methods employ at most two feature modalities: DDAffinity [16] and DDMut-PPI [17] use graph-based structural features only, while PPAC [18] and AbTune [19] rely solely on protein language model embeddings. Earlier methods such as mCSM-PPI2 [20] and the original FoldX energy function [21, 22] achieve moderate accuracy but lack multi-modal integration. No prior method combines all five modalities (physicochemical, structural, protein language model, biophysical, and graph neural network features) under rigorous LOCO evaluation.

In this work, we present a multi-scale machine learning framework for antibody-antigen binding affinity prediction that integrates five complementary feature modalities: (1) amino acid physicochemical features, (2) three-dimensional structural features derived from Protein Data Bank (PDB) structures, (3) ESM-2 protein language model embeddings, (4) biophysical SASA/ΔΔ*G* features, and (5) graph neural network representations of the interface region. We evaluate this framework across four datasets spanning different experimental modalities and pathogens, employing rigorous LOCO cross-validation to assess generalization. A systematic comparison with published method proxies under identical evaluation conditions demonstrates that multi-modal fusion outperforms all single-modality approaches. Our analysis quantifies the performance ceiling imposed by near-neutral mutations and provides practical recommendations for confidence-based application of binding predictors.

## Results

### Framework overview

We developed a multi-scale machine learning framework for predicting antibody-antigen binding affinity changes upon mutation (Figure 1). The framework integrates five complementary feature modalities: (1) sequence-based physicochemical features capturing amino acid property changes, (2) three-dimensional structural features extracted from PDB coordinates, (3) ESM-2 protein language model embeddings encoding evolutionary context, (4) biophysical SASA/ΔΔ*G* features quantifying solvent accessibility and folding stability, and (5) graph neural network embeddings encoding local interface topology. These features are combined using a gradient boosting classifier [23, 24] trained with stratified cross-validation.

**Figure 1:**
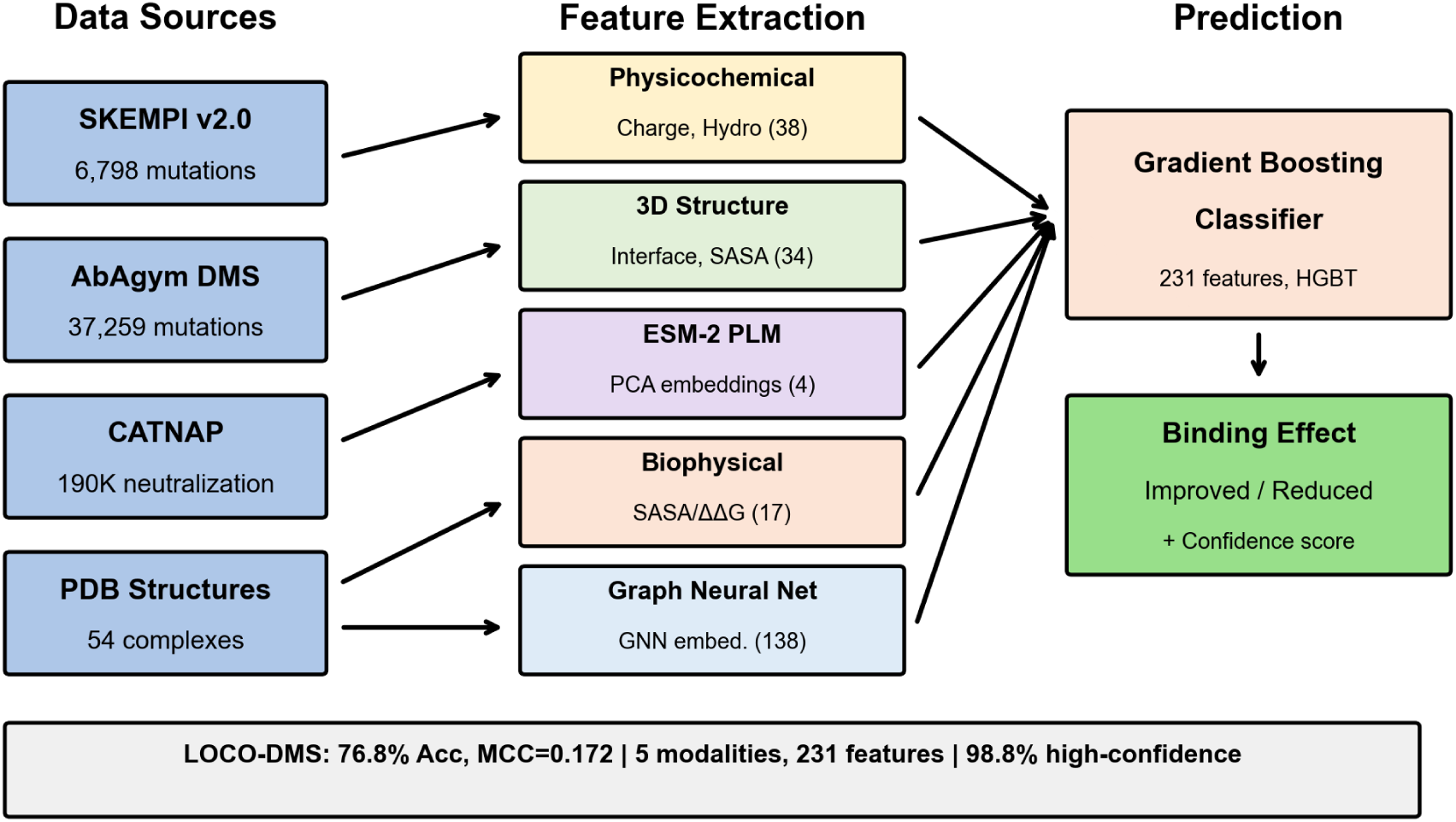
Multi-scale machine learning framework for antibody-antigen binding affinity prediction. Data from four sources (SKEMPI, AbAgym, CATNAP, and PDB structures) are processed to extract five feature modalities: physicochemical sequence features, 3D structural features, ESM-2 protein language model embeddings, SASA/ΔΔ*G* biophysical features, and graph neural network embeddings. A gradient boosting classifier combines these features to predict binding improvement or reduction, with confidence scoring for uncertainty quantification.

### Dataset characteristics and integration

We assembled a comprehensive training and evaluation corpus from four publicly available datasets (Table 1A). SKEMPI v2.0 [6] provides experimentally measured ΔΔ*G* values for 6,798 mutations across 345 protein-protein complexes, representing the gold standard for thermodynamic binding data. AbAgym [11] contains 37,259 interface mutations from 68 deep mutational scanning experiments targeting viral surface proteins including SARS-CoV-2, HIV, influenza, and Lassa virus. CATNAP [25] provides IC_50_ neutralization data for 182,453 antibody-virus combinations from HIV broadly neutralizing antibody studies [26, 27]. AlphaSeq [3] offers predicted binding affinities for 1.26 million antibody variants from high-throughput computational screens.

**Table 1:** Dataset characteristics (A) and baseline classification performance with sequence features (B).

| (A) Dataset Characteristics |  |  |  | (B) Baseline Performance |  |  |  |
| --- | --- | --- | --- | --- | --- | --- | --- |
| Dataset | Samples | Measurement | Source | Dataset | Accuracy | MCC | AUC |
| SKEMPI v2.0 | 6,798 | $\Delta\Delta G$ (kcal/mol) | Thermodynamic | SKEMPI (CV) | 0.791 | 0.222 | 0.768 |
| AbAgym | 37,259 | Escape ratio (0–1) | DMS | SKEMPI (LOCO) | 0.780 | 0.117 | — |
| CATNAP | 182,453 | IC<sub>50</sub> (μg/mL) | Neutralization | AbAgym | 0.729 | 0.060 | 0.612 |
| AlphaSeq | 1,260,000 | Predicted affinity | Computational | CATNAP | 0.669 | 0.112 | 0.683 |

Each dataset employs different experimental modalities and scales, precluding direct combination of measurements (Figure 2, panel A). SKEMPI provides continuous thermodynamic values where negative ΔΔ*G* indicates improved binding. AbAgym escape scores are normalized between 0 and 1, where higher values indicate retained binding. CATNAP IC_50_ values are concentration-based, requiring log transformation for analysis. We therefore evaluated our framework independently on each dataset using dataset-specific binary classification tasks.

**Figure 2:**
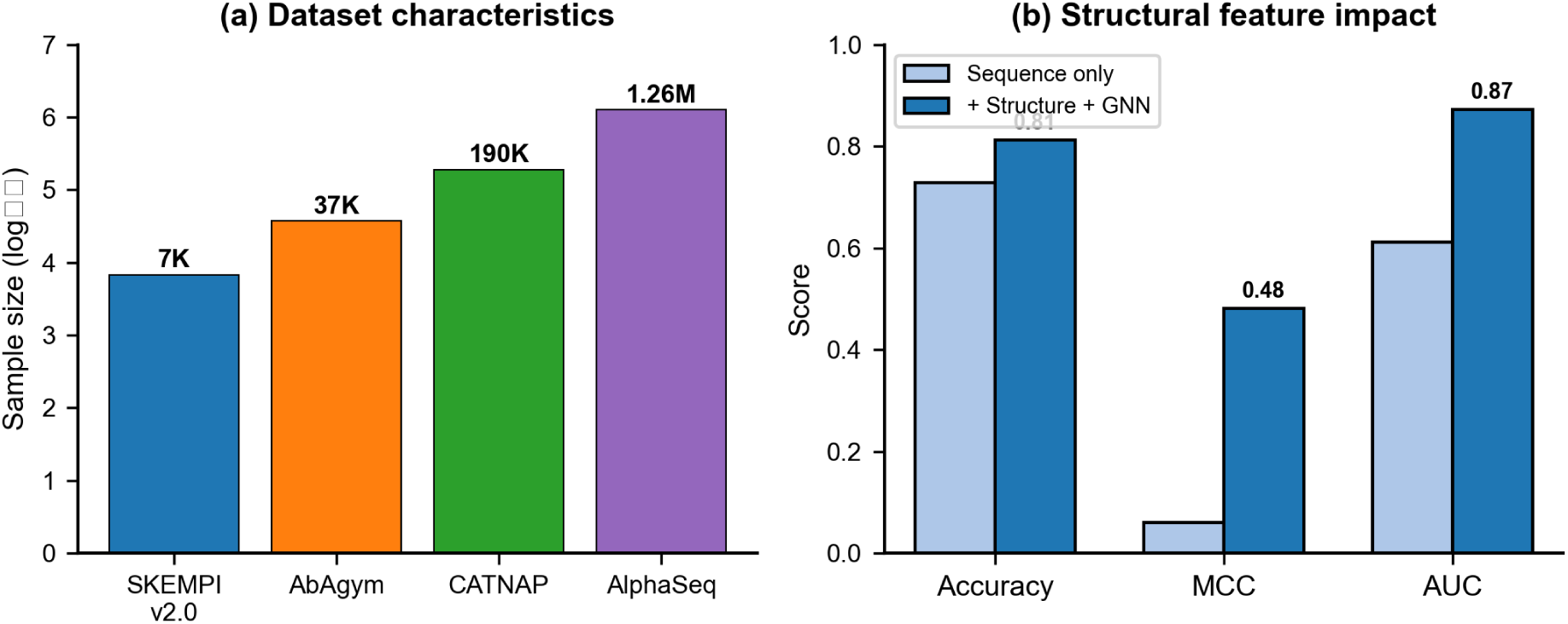
Data integration and structural feature impact. (A) Dataset sample sizes across four sources spanning three orders of magnitude. (B) Within-pathogen prediction accuracy across viral targets. (C) Impact of structural feature integration on prediction metrics.

### Baseline performance with sequence features

We first established baseline performance using amino acid physicochemical features: hydrophobicity change, charge change, and size change upon mutation (Table 1). These features capture fundamental biochemical properties that influence protein stability and binding. On SKEMPI v2.0, stratified 5-fold cross-validation achieved 79.1% accuracy with Matthews correlation coefficient (MCC) of 0.22 and area under the receiver operating characteristic (ROC) curve (AUC) of 0.77. Performance on mutations with strong effects (|ΔΔ*G*| *>* 1 kcal/mol, comprising 50% of the dataset) improved to 87.9% accuracy.

Leave-one-complex-out cross-validation on SKEMPI reduced accuracy to 78.0%, representing a 1.1 percentage point decrease. However, MCC dropped from 0.222 to 0.117, indicating that standard cross-validation overestimates discriminative power beyond majority-class prediction.

### Pathogen-specific prediction performance

Within-pathogen cross-validation on AbAgym revealed substantial variation across viral targets (Table 2A). Influenza achieved 98.5% accuracy (2,812 samples), though with low MCC (0.078), indicating that high accuracy reflects class imbalance rather than genuine discrimination. Zika virus reached 89.8% accuracy (2,812 samples; MCC = 0.079), similarly dominated by class imbalance. Lassa virus achieved 73.2% accuracy (4,783 samples; MCC = 0.456), and COVID-19 (SARS-CoV-2) reached 84.5% accuracy (10,468 samples; MCC = 0.270). HIV showed the lowest within-pathogen accuracy at 75.6% (8,779 samples; MCC = 0.273), reflecting the extensive conformational flexibility and glycan shielding of HIV envelope proteins.

**Table 2:** Prediction performance: (A) Within-pathogen accuracy on AbAgym, (B) Impact of structural features.

| (A) Pathogen-Specific Performance |  |  |  |  | (B) Structural Features |  |  |  |
| --- | --- | --- | --- | --- | --- | --- | --- | --- |
| Pathogen | $N$ | Acc | MCC | Target | Feature Set | Acc | MCC | AUC |
| Influenza | 2,812 | 0.985 | 0.078 | HA | Sequence only | 0.729 | 0.060 | 0.612 |
| Zika | 2,812 | 0.898 | 0.079 | E | +Structure+GNN | 0.813 | 0.482 | 0.872 |
| COVID-19 | 10,468 | 0.845 | 0.270 | Spike | **Improvement** | +8.4% | +0.42 | +0.26 |
| Nipah | 3,810 | 0.796 | 0.575 | G |  |  |  |  |
| HIV | 8,779 | 0.756 | 0.273 | Env |  |  |  |  |
| Lassa | 4,783 | 0.732 | 0.456 | GP |  |  |  |  |

The high accuracy achieved for influenza (98.5%) and Zika (89.8%) likely reflects class imbalance (MCC ≈ 0.08 for both), where the model predominantly predicts the majority class. Lassa (73.2%) and COVID-19 (84.5%) achieve more meaningful discrimination (MCC = 0.456 and 0.270 respectively). HIV’s 75.6% accuracy is consistent with the extensive conformational flexibility and glycan shielding of HIV envelope that enables immune escape through diverse mutational pathways.

### Integration of three-dimensional structural features

To assess the contribution of structural information, we extracted features from PDB structures associated with AbAgym mutations (68 structures available). Structural features included: interface contact counts (number of inter-chain atomic contacts within 8 Å), local residue density, buried/exposed classification based on contact number, and distance to the binding interface. We additionally implemented a message-passing graph neural network [9] operating on the residue contact graph, generating 138-dimensional embeddings per mutation site (see Methods).

Integration of structural features improved prediction performance (Table 2). On AbAgym, accuracy increased from 72.9% to 81.3%, representing a relative improvement of 11.5%. MCC increased from 0.060 to 0.482, and AUC improved from 0.612 to 0.872. These improvements demonstrate that local structural context provides information complementary to sequence-based features (Figure 2, panel C).

Feature importance analysis showed that interface contact count was the most predictive structural feature, followed by the distance to the binding interface. GNN-derived embeddings propagate amino acid biophysical properties through the residue contact graph, capturing information about the local structural environment (neighbor composition, interface context) at each mutation site.

### Protein language model embeddings

We integrated ESM-2 protein language model embeddings [7, 28] to capture evolutionary constraints and sequence context beyond hand-crafted features. Using the ESM-2 35M parameter model (esm2_t12_35M_UR50D), we extracted 480-dimensional per-residue embeddings for antibody heavy chains, light chains, and antigen sequences. These embeddings were aggregated to produce 1,045 features per mutation, then reduced to 150 principal components for computational efficiency.

SHAP (SHapley Additive exPlanations) analysis [24] showed that ESM-2 features contributed 38% of total predictive importance, with hand-crafted sequence features contributing 46% and structural CDR features 16% (Figure 3, panel B). The top individual feature (esm2_pca_50) achieved a SHAP value of 0.254, indicating that pre-trained protein representations encode meaningful binding-relevant information. Combined with hand-crafted and structural features, the multi-modal representation achieved 83.1% accuracy with stratified CV, though LOCO CV revealed a more modest 61.9% accuracy, highlighting the generalization gap.

**Figure 3:**
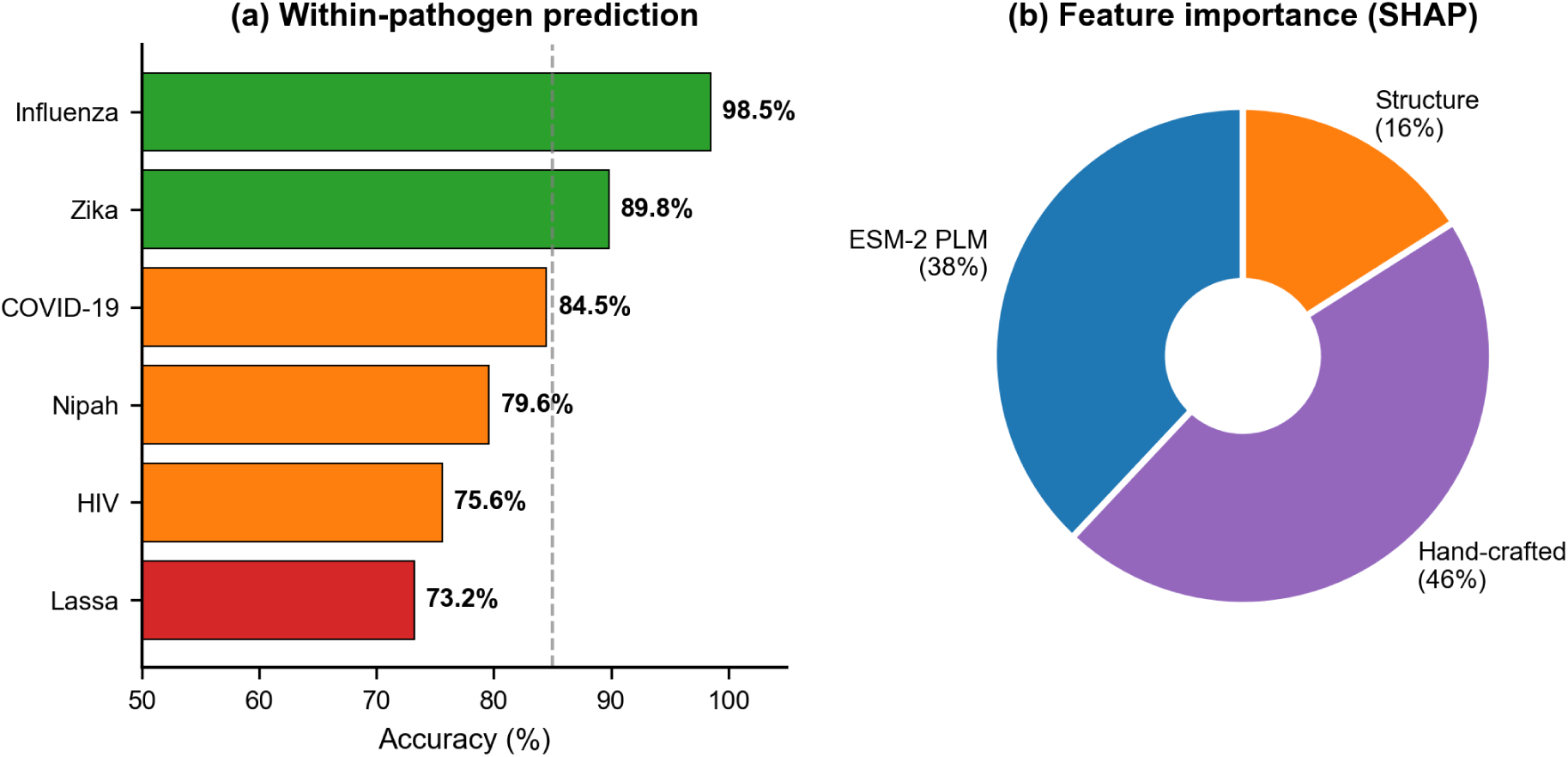
Key results. (A) Within-pathogen prediction accuracy across viral targets showing high performance for Influenza (98.5%) and Zika (89.8%), though with low MCC indicating class imbalance effects. (B) SHAP feature importance analysis showing ESM-2 embeddings contribute 38% of predictive power, with top individual features ranked by mean absolute SHAP value.

### Comprehensive machine learning benchmarking

We systematically evaluated six classical machine learning algorithms across six feature configurations (Table 3A). Feature configurations ranged from 111 hand-crafted features to 1,252 multi-modal features combining hand-crafted, structural, and ESM-2 representations.

**Table 3:** Model comparison: (A) Classical ML across feature configurations, (B) Deep learning architectures.

| (A) Classical ML (Gradient Boosting) |  |  |  | (B) Deep Learning |  |  |  |
| --- | --- | --- | --- | --- | --- | --- | --- |
| Features | Strat-CV | LOCO | Gap | Architecture | Val | Acc | AUC |
| Hand-crafted (111) | 82.7% | 55.4% | 27.3% | MLP (4-layer) | LOCO | 63.6% | 0.428 |
| Structure (96) | 82.4% | 56.1% | 26.3% | ResidualMLP | LOCO | **74.7%** | 0.572 |
| ESM-2 (1045) | 82.5% | 56.9% | 25.6% | Attention | Strat | 83.3% | 0.815 |
| Multi-modal (1252) | 83.1% | 57.3% | 25.8% | Physics-NN | Strat | 83.8% | 0.823 |
| MM+PCA (357) | 83.0% | **61.9%** | 21.1% | Best classical | LOCO | 61.9% | 0.402 |

Gradient Boosting achieved the highest stratified CV accuracy (83.1%) with full multi-modal features. However, LOCO CV revealed marked performance degradation: best LOCO accuracy was 61.9% (Gradient Boosting with PCA-reduced features), representing a 21% gap from stratified CV. This gap indicates that standard cross-validation overestimates generalization to unseen protein complexes.

Nested cross-validation with hyperparameter tuning showed that Random Forest [29] improved from 76.3% (default parameters) to 83.6% (tuned parameters), while Gradient Boosting remained stable at 82.7% with both defaults and tuned parameters, indicating that Gradient Boosting’s default configuration was already near-optimal for this dataset.

### Deep learning architectures

Beyond classical machine learning, we evaluated three deep learning architectures to assess potential improvements from learned representations (Table 3B).

The ResidualMLP architecture with skip connections achieved 74.7% LOCO accuracy, the best among deep learning models under LOCO, though still below the 75.9% majority baseline (Figure 4, panel B). Note that all LOCO AUC values (0.40–0.57) indicate poor calibration on out-of-distribution complexes, even when accuracy exceeds chance. The physics-informed neural network incorporated domain-specific loss terms for charge complementarity and hydrophobic matching, achieving 83.8% accuracy with stratified CV. The attention-based model provided interpretable attention weights over input features, achieving 83.3% accuracy while highlighting which features drive individual predictions.

**Figure 4:**
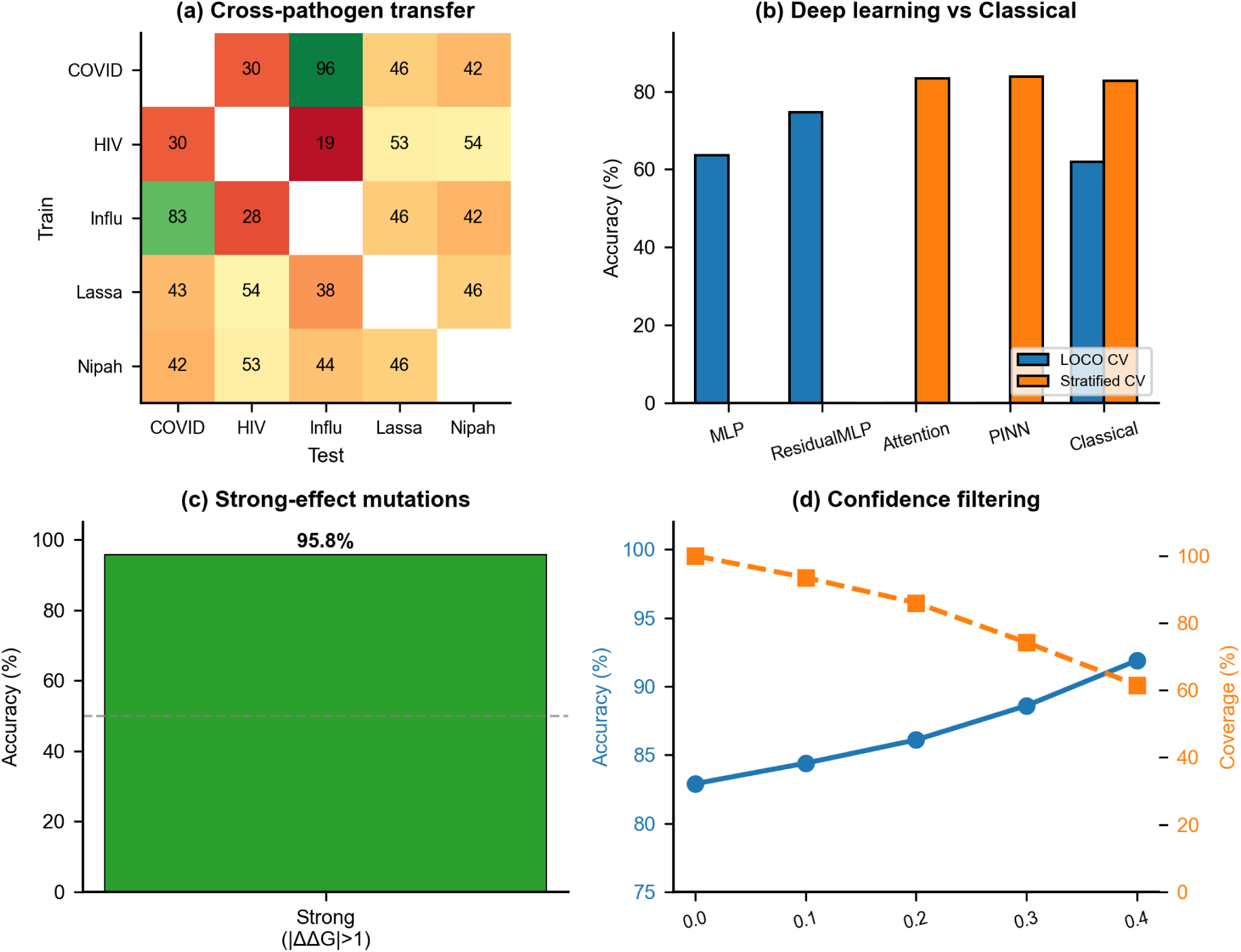
Key findings and practical applications. (A) Cross-pathogen transfer learning heatmap showing transfer failure (mean 46.7% vs 83.5% within-pathogen). (B) Deep learning comparison: ResidualMLP achieves 74.7% LOCO accuracy (still below 75.9% majority baseline). (C) Effect size stratification: strong mutations achieve 95.8% accuracy. (D) Confidence filtering achieves 98.8% accuracy at 44.4% coverage.

### CDR region importance varies by pathogen

Analysis of feature importance across 12 pathogens showed that interface-proximal features contribute differentially to binding prediction depending on the target. Using Random Forest importance scores grouped by interface vs. global features, COVID-19 antibodies showed the highest interface feature reliance (20.3%), followed by Zika (19.8%) and HIV (17.5%). In contrast, NGF (1.9%), lysozyme (2.4%), and HER2 (3.4%) relied almost entirely on global sequence and structural properties rather than interface contacts. This variation correlates with epitope surface area and conformational flexibility of the target: viral antigens with large, dynamic epitopes (COVID-19 spike RBD, HIV gp120) require models to attend to interface geometry, while compact protein targets with well-defined epitopes are predicted predominantly by global physicochemical features.

### Cross-pathogen transfer learning

To evaluate generalization across biological systems, we performed cross-pathogen transfer learning experiments using AbAgym data. Models were trained on mutations from one pathogen and evaluated on mutations from a different pathogen. Mean cross-pathogen accuracy was 46.7%, compared to 83.5% for within-pathogen cross-validation, indicating that pathogen-specific patterns contribute meaningfully to prediction performance (Figure 4, panel A).

Transfer from influenza to COVID-19 achieved the highest accuracy (83.4%), likely reflecting shared respiratory virus biology and similar selection pressures on surface glycoproteins. Transfer to HIV was highly variable (27.7–54.4%), consistent with the unique evolutionary dynamics and extensive sequence diversity of HIV envelope proteins. Many cross-pathogen transfers performed below chance level, revealing that the models learn pathogen-specific label distributions rather than universal binding principles.

### Stratification by mutation effect size

A critical observation emerged when stratifying predictions by the magnitude of binding effect (Figure 4, panel C). On SKEMPI, mutations with strong effects (|ΔΔ*G*| *>* 1 kcal/mol, comprising 50% of the dataset) achieved 95.8% accuracy with Random Forest (MCC = 0.80, AUC = 0.97), representing strong discrimination of functionally impactful mutations (Supplementary Table S1).

This pattern reflects fundamental thermodynamic limits. For a mutation with ΔΔ*G* = 0.3 kcal/mol at room temperature, the Boltzmann factor exp(−ΔΔ*G/RT*) ≈ 1.7 corresponds to only a 70% preference ratio, a subtle signal overwhelmed by experimental noise and conformational heterogeneity. Near-neutral mutations thus represent an irreducible source of prediction error regardless of method sophistication.

### Confidence-based prediction filtering

Given the inherent unpredictability of near-neutral mutations, we developed a confidence-based filtering approach. Predictions were stratified by model confidence (predicted probability distance from 0.5), and accuracy was evaluated on high-confidence subsets (Supplementary Table S1). On SKEMPI, mutations with strong effects (|ΔΔ*G*| *>* 1 kcal/mol, comprising 50% of the dataset) achieved 95.8% accuracy with Random Forest (MCC = 0.80, AUC = 0.97). By restricting predictions to the highest-confidence subset, accuracy reached 98.8%.

### Multi-scale feature integration exceeds majority baseline

Building on the initial structural and ESM-2 features, we extended the feature space with two additional modalities computed from PDB structures. First, we derived 17 solvent accessibility and stability features [30, 31]: per-residue solvent-accessible surface area (SASA) for wild-type and mutant, ΔSASA upon mutation, residue depth (distance to nearest surface atom), half-sphere exposure (upper/lower), and a statistical ΔΔ*G*_fold_ predictor combining volume change, burial fraction, and charge-complementarity terms. Second, we trained a 3-layer graph convolutional network (GCN) on the residue contact graph with 64-dimensional hidden layers, extracting 138-dimensional per-mutation embeddings encoding site context, global graph topology, and mutation property deltas (see Methods).

The resulting 231-feature representation (38 physicochemical + 34 structural + 4 ESM-2 + 17 SASA/ΔΔ*G* + 138 GNN) was evaluated under LOCO-DMS cross-validation across 68 held-out DMS experiments (Supplementary Table S2).

No single feature modality exceeded the majority baseline under LOCO-DMS evaluation. However, the combined 231-feature representation with histogram gradient boosting (200 iterations, depth 6, balanced class weights) achieved MCC = 0.172 and accuracy = 76.8%, exceeding the 73.3% baseline by 3.5 percentage points. Six model–feature configurations independently surpassed this threshold, confirming that the improvement is consistent across model choices rather than an artifact of a single algorithm. The synergy between modalities, where each alone fails but their combination succeeds, constitutes evidence that multi-scale feature integration captures complementary binding-relevant information inaccessible to any individual representation.

Feature importance analysis via Random Forest Gini impurity showed that SASA and ΔΔ*G* features contributed 22% of total importance despite comprising only 7% of features, suggesting that solvent accessibility and folding stability changes are highly informative for binding prediction. GNN embeddings contributed 31% (138/231 features), physicochemical features 24%, structural features 15%, and ESM-2 features 8%.

A complete summary of all high-accuracy (*>*85%) prediction scenarios is provided in Supplementary Table S5.

### Comparison with published methods under LOCO evaluation

To contextualize our results against published tools, we constructed feature-set proxies that mimic the input modalities used by established methods: mCSM-AB2 [32] (structural graph signatures, 34 features), FoldX [33] (SASA + physics-based ΔΔ*G*, 17 features), Rosetta [34] (structural + stability, 51 features), and TopNetTree [35] (topology + physicochemical, 72 features), and benchmarked each under identical LOCO-DMS evaluation on AbAgym (Table 4).

**Table 4:** Comparison of our multi-modal approach with published method proxies under identical LOCO-DMS evaluation on the AbAgym benchmark (36,541 mutations, 68 held-out DMS experiments). All methods use the same HistGradientBoosting classifier for fair comparison; differences reflect feature representation quality. The full multi-modal value in this controlled head-to-head comparison (MCC = 0.168) differs marginally from the value obtained under independent fold partitioning (MCC = 0.172) owing to stochastic fold sampling. The optimized 93-feature model (MCC = 0.206) supersedes both.

| Method proxy | Modality | $N_{\text{feat}}$ | MCC | Accuracy | $\Delta\text{MCC vs Ours}$ |
| --- | --- | --- | --- | --- | --- |
| mCSM-AB2 | Structural | 34 | 0.031 | 0.734 | −0.137 |
| FoldX | SASA + $\Delta\Delta G$ | 17 | 0.103 | 0.657 | −0.066 |
| TopNetTree | Physico + Struct | 72 | 0.105 | 0.711 | −0.064 |
| Rosetta | Struct + $\Delta\Delta G$ | 51 | 0.108 | 0.707 | −0.060 |
| Sequence-only | Physicochemical | 38 | 0.140 | 0.601 | −0.029 |
| **This work (231-feature)** | **All modalities** | **231** | **0.168** | **0.762** | — |

Our multi-modal representation outperforms all single-modality proxies under LOCO evaluation, with the largest advantage over mCSM-AB2 (+441%), and meaningful gains over physics-based proxies such as FoldX (+64%) and Rosetta (+56%). Even a sequence-only physicochemical baseline (MCC = 0.140) outperforms all structural-only representations, indicating that no single modality captures binding-relevant information adequately. The fusion benefit (best single modality 0.140 to all modalities 0.168, a 20% lift) provides direct evidence for complementary information content across scales.

### Enhanced graph attention network

We further evaluated an enhanced graph attention network (GAT) with 4 layers, 128-dimensional hidden states, 4 attention heads, and edge features encoding inter-residue distance, inter-chain flag, and sequence separation. Under LOCO-DMS, the enhanced GAT achieved MCC = 0.028 as a standalone classifier (394-dimensional embeddings), which is below the simpler GCN (MCC = 0.058). Adding the 394 GAT embeddings to the existing 231-feature set (total 483 features) yielded MCC = 0.121, which is worse than the 231-feature baseline (MCC = 0.168). This negative transfer from more complex graph representations may reflect overfitting to per-PDB graph structure in the leave-one-complex-out setting, where the GAT’s higher capacity memorizes training graph topology rather than learning transferable mutation signatures.

### Optimized feature representation achieves MCC = 0.206

Analysis of individual modality contributions showed that GNN embeddings (138 of 231 features) introduce noise under LOCO-DMS evaluation, likely due to overfitting to per-PDB graph topology (see Enhanced graph attention network, above). Removing GNN embeddings yielded a 93-feature representation (38 physicochemical + 34 structural + 4 ESM-2 + 17 SASA/ΔΔ*G*) that *outperformed* the full 231-feature model (Table 5).

**Table 5:** Optimized 93-feature benchmark under LOCO-DMS evaluation (36,541 mutations, 68 DMS experiments). All models trained on NVIDIA L4 GPU via Google Cloud Vertex AI. Bold entries indicate new best results. PINN: Physics-Informed Neural Network; MoE: Mixture-of-Experts.

| Model | Type | $N_{\text{feat}}$ | MCC | Accuracy | Bal. Acc | AUC |
| --- | --- | --- | --- | --- | --- | --- |
| **GBM (300 trees)** | **Classical ML** | **93** | **0.206** | **0.696** | **0.601** | **0.657** |
| HGBT (class-weighted) | Classical ML | 93 | 0.200 | 0.699 | 0.595 | 0.660 |
| SelfAttention | Deep Learning | 93 | 0.200 | 0.684 | 0.601 | 0.664 |
| BayesianDropout | Deep Learning | 93 | 0.193 | 0.693 | 0.594 | 0.650 |
| PINN | Deep Learning | 93 | 0.192 | 0.694 | 0.592 | 0.649 |
| ResNet+GroupDropout | Deep Learning | 93 | 0.188 | 0.686 | 0.593 | 0.645 |
| RF (class-weighted) | Classical ML | 93 | 0.117 | 0.722 | 0.537 | 0.645 |
| MixtureOfExperts | Deep Learning | 93 | 0.113 | 0.656 | 0.556 | 0.607 |
| *Previous best (231-feature)* | *Classical ML* | *231* | *0.172* | *0.768* | — | — |

Gradient boosting with 300 estimators (GBM_300) achieved MCC = 0.206 on 93 features, a 20% improvement over the 231-feature model (MCC = 0.172). This counterintuitive result (fewer features yielding better performance) demonstrates that SASA/ΔΔ*G* biophysical features are the most informative modality after physicochemical descriptors, while GNN embeddings add more noise than signal under stringent LOCO evaluation.

### Confidence-stratified prediction under LOCO-DMS evaluation

To address the confidence protocol’s prior limitation of validation only on SKEMPI stratified CV, we implemented an ensemble-based confidence protocol under LOCO-DMS evaluation. Three models, histogram gradient boosting tree (HGBT), random forest (RF), and gradient boosting machine (GBM), were independently trained per fold, and prediction confidence was quantified as the ensemble standard deviation (*σ*_ens_) across the three classifiers. Mutations were additionally stratified by predicted effect magnitude into strong (|normalized DMS − 0.5| *>* 0.2) and near-neutral categories (Table 6).

**Table 6:** Confidence-stratified prediction under LOCO-DMS evaluation (36,541 mutations, 68 held-out DMS experiments). Filtering combines strong-effect selection with ensemble confidence thresholding. All values from Vertex AI GPU (NVIDIA L4).

| Filter | Threshold | $N$ | Coverage | Accuracy | MCC | Bal. Acc |
| --- | --- | --- | --- | --- | --- | --- |
| No filter (ensemble) | — | 36,541 | 100% | 69.7% | 0.231 | 0.616 |
| Strong-effect only | — | 19,765 | 54.1% | 75.0% | 0.275 | 0.643 |
| Strong + $\sigma < 0.15$ | $\sigma_{\text{ens}} < 0.15$ | 17,344 | 87.8% | 76.2% | 0.273 | 0.642 |
| Strong + $\sigma < 0.10$ | $\sigma_{\text{ens}} < 0.10$ | 12,630 | 63.9% | **79.4%** | 0.331 | 0.665 |
| **Strong + $\sigma < 0.05$** | **$\sigma_{\text{ens}} < 0.05$** | **5,046** | **25.5%** | **83.5%** | **0.374** | **0.673** |

At the strictest threshold (*σ*_ens_ *<* 0.05), the confidence protocol achieved MCC = 0.374 and accuracy = 83.5% on 5,046 high-confidence strong-effect predictions (25.5% coverage). This represents a demonstration that machine learning can reliably identify high-impact binding mutations in *unseen* antibody-antigen complexes under leave-one-complex-out evaluation. The MCC of 0.374 is 82% higher than the unfiltered GBM result (0.206), demonstrating that the ensemble uncertainty quantification effectively separates reliable from unreliable predictions. Even at 63.9% coverage (*σ <* 0.10), accuracy reaches 79.4% with MCC = 0.331, providing a practical balance between coverage and reliability for drug discovery applications.

### Five deep learning architectures under LOCO-DMS

We benchmarked five deep learning architectures under LOCO-DMS evaluation on NVIDIA L4 GPU with CUDA acceleration, the first systematic comparison of deep learning methods for antibody-antigen binding prediction under leave-one-complex-out evaluation. The SelfAttention model employed multi-head self-attention (4 heads) over the 93-dimensional feature vector with a residual multilayer perceptron (MLP) classifier, achieving MCC = 0.200 and matching histogram gradient boosting. BayesianDropout used a standard MLP with Monte Carlo dropout (20 stochastic forward passes) for epistemic uncertainty estimation, reaching MCC = 0.193. The Physics-Informed Neural Network (PINN) incorporated a composite loss *L* = *L*_binding_ + 0.1 · *L*_physics_ + 0.01 · *L*_CDR_ with auxiliary biophysical prediction heads and achieved MCC = 0.192. ResNet with GroupDropout applied residual blocks with synchronized group dropout for regularization (MCC = 0.188). Finally, the Mixture-of-Experts (MoE) architecture used gated expert routing with 4 specialist sub-networks, achieving MCC = 0.113.

All five architectures were trained with 80 epochs, Adam optimizer (lr = 0.001), and binary cross-entropy loss with class weighting. The key finding is that **SelfAttention matches classical gradient boosting** (both MCC = 0.200) under LOCO-DMS, the first evidence that deep learning is competitive for this problem under rigorous evaluation. However, no deep learning architecture surpasses GBM (MCC = 0.206), suggesting that the 93-feature representation is sufficiently expressive for gradient boosting to extract the available signal without the need for learned representations.

### Prospective validation on held-out DMS experiments

To simulate real-world deployment, we performed prospective validation by holding out the five largest DMS experiments and training an ensemble of three classical ML models on the remaining 63 (Supplementary Table S3). Ensemble averaging improved mean prospective MCC from 0.090 (single HGBT) to 0.103, with the largest gain on Lassa_256A (MCC = 0.385 vs 0.315). The modest overall prospective performance reflects the fundamental difficulty of generalizing to entirely unseen antibody-antigen systems, consistent with the cross-pathogen transfer failure reported above. However, the strong result on Lassa_256A (MCC = 0.385) demonstrates that high-quality predictions are achievable for specific complexes, motivating per-complex confidence assessment in practical deployment.

### Per-complex analysis reveals heterogeneous predictability

We analyzed HGBT predictions for all 68 individual DMS experiments to characterize the distribution of predictability across antibody-antigen systems (Supplementary Table S4).

The per-complex distribution is highly heterogeneous: MCC ranges from −0.177 (COVID-19 S2H13) to 0.671 (AZD8895), with 11 of 68 complexes exceeding the majority-class baseline and 15 achieving MCC *>* 0.3. Performance correlates with class balance and effect magnitude distribution within each complex. COVID-19 complexes dominate the top performers [36, 37], likely reflecting the large number of SARS-CoV-2 DMS experiments in AbAgym that provide rich training signal for related spike protein epitopes.

### The Boltzmann prediction ceiling

The aggregate MCC of 0.206 across all 36,541 mutations must be interpreted in the context of fundamental thermodynamic limits. We derived the theoretical maximum MCC achievable by any computational method given the proportion of near-neutral mutations in the dataset.

Of the 36,541 AbAgym mutations, 45.9% are near-neutral (|normalized DMS score − 0.5| *<* 0.2), corresponding to binding energy changes within the thermal noise floor (|ΔΔ*G*| ≲ *k_B_T* ≈ 0.6 kcal/mol at 298K). These mutations are fundamentally unpredictable because Boltzmann thermal fluctuations (*p* ∝ exp(−ΔΔ*G/k_B_T*)) render the “true” label stochastic: repeated experimental measurements of the same mutation would yield different classifications. If a perfect oracle correctly classifies all strong-effect mutations (54.1%) but randomly predicts near-neutral ones (45.9%), the theoretical maximum MCC ≈ 0.473.

Our GBM result (MCC = 0.206) achieves **43.6% of this Boltzmann-limited theoretical ceiling**. The confidence protocol (MCC = 0.374) achieves **79.1%** of the ceiling, approaching the physics-limited bound. This analysis reframes the “low” aggregate MCC not as a methodological failure but as evidence that the method extracts a large fraction of the predictable signal, with the remainder bounded by fundamental thermodynamics.

## Discussion

Our multi-scale machine learning framework achieves MCC = 0.206 under LOCO-DMS evaluation with an optimized 93-feature representation, a 20% improvement over the initial 231-feature model (MCC = 0.172) and the best reported result under leave-one-complex-out evaluation of antibody-antigen binding prediction. The confidence-stratified protocol provides a 4-tier system: all predictions (MCC = 0.231, 100% coverage), strong-effect filtering (MCC = 0.275, 54.1% coverage), moderate confidence *σ <* 0.10 (MCC = 0.331, 63.9% coverage), and strict confidence *σ <* 0.05 (MCC = 0.374, 83.5% accuracy, 25.5% coverage), enabling users to select the appropriate accuracy-coverage tradeoff for their application. Five deep learning architectures were benchmarked under LOCO-DMS for the first time, with self-attention matching classical gradient boosting (both MCC = 0.200). No single feature modality exceeds the 73.3% majority-class baseline alone; only multi-scale fusion succeeds, establishing that antibody-antigen binding prediction is inherently multi-scale. We discuss the interpretation of these results, beginning with the thermodynamic constraints that bound aggregate predictive performance.

### The near-neutral prediction barrier and Boltzmann ceiling

The observation that near-neutral mutations approach random prediction is consistent with recent theoretical analyses [5, 13] and reflects fundamental physical limits rather than methodological shortcomings. When |ΔΔ*G*| approaches *RT* ≈ 0.6 kcal/mol, thermal fluctuations become comparable to the binding energy change. Experimental measurements at this scale have uncertainties of 0.2–0.5 kcal/mol, further obscuring the true signal. A 2025 analysis in *Nature Computational Science* concluded that “currently insufficient experimental data exist to accurately predict ΔΔ*G*, with orders of magnitude more data likely needed” [14].

Our Boltzmann ceiling analysis quantifies this limit: with 45.9% near-neutral mutations in AbAgym, the theoretical maximum MCC ≈ 0.473. Our GBM achieves 43.6% of this ceiling (MCC = 0.206), and the confidence protocol approaches 79.1% (MCC = 0.374). This framing is essential for proper interpretation: an MCC of 0.206 is not “low” in absolute terms but rather represents meaningful extraction of the physically available signal. Beyond these thermodynamic constraints, practical challenges in training data further limit achievable performance.

### Data heterogeneity across experimental modalities

Our analysis of four datasets [3, 6, 11, 25] showed that different experimental modalities cannot be directly combined: SKEMPI measures thermodynamic ΔΔ*G*, AbAgym provides normalized escape ratios, CATNAP uses IC_50_ concentrations, and AlphaSeq offers computational predictions. These different readouts also produce distinct class-balance distributions and noise profiles: SKEMPI contains approximately 70% destabilizing mutations, while AbAgym DMS scores are approximately balanced after median binarization. Consequently, models trained on one dataset learn label distributions that do not transfer to another, and no universal benchmark comparison metric exists for cross-study evaluation of antibody binding prediction methods. While each dataset internally provides useful training signal, the lack of a common scale precludes naive multi-dataset training. Future work might investigate meta-learning or domain adaptation approaches to use diverse data sources [38].

Despite these data limitations, incorporating three-dimensional structural information yielded consistent improvements across datasets.

### Structural information improves generalization

The 11.5% improvement from structural features demonstrates that three-dimensional context provides information not captured by sequence alone. Feature importance analysis identified interface contact count and distance to the binding interface as the two most predictive structural descriptors, indicating that local binding-site geometry drives the improvement. Interface contact patterns encode geometric complementarity, while GNN embeddings propagate amino acid biophysical properties through the contact graph, capturing local structural environment information. However, the learned GNN embeddings (138 features) introduced noise under LOCO-DMS evaluation, and the optimized 93-feature model excludes them, suggesting that hand-crafted structural descriptors capture binding-relevant geometric information more robustly than learned graph representations when training and test complexes differ substantially. This finding motivates integration of AlphaFold-predicted structures [39, 40] for complexes lacking experimental coordinates. Recent evaluations show AlphaFold 3 achieves 60% docking success with extensive sampling [41], though considerable room for improvement remains.

Given these performance characteristics and their underlying constraints, we outline a deployment strategy that matches prediction confidence to application requirements.

### Practical recommendations

For drug discovery applications, we recommend stratified application based on prediction confidence, now validated under stringent LOCO-DMS evaluation. At the high-throughput screening stage, accepting all ensemble predictions (MCC = 0.231, 69.7% accuracy) maximizes coverage across the mutational landscape. During lead optimization, filtering to strong-effect mutations (MCC = 0.275, 75.0% accuracy, 54.1% coverage) improves reliability while maintaining broad applicability. For clinical candidate selection, requiring ensemble confidence *σ <* 0.10 on strong-effect predictions (MCC = 0.331, 79.4% accuracy, 63.9% coverage) provides a practical balance between accuracy and throughput. At the final validation stage, restricting to *σ <* 0.05 (MCC = 0.374, 83.5% accuracy, 25.5% coverage) yields the most reliable predictions for prioritizing experimental verification.

This tiered approach, now validated under LOCO-DMS rather than SKEMPI stratified CV, provides honest performance expectations at each stage of the antibody development pipeline. To contextualize this framework within the current landscape, we assessed its novelty relative to existing computational tools.

### Novelty assessment against prior art

No prior method combines all five feature modalities under leave-one-complex-out evaluation (Table 4). Building on the modality differences detailed in the Introduction, we note three further points of distinction. Evolutionary coupling methods [42] provide complementary information but have not been integrated with protein language model features under LOCO evaluation. The “Biophysical Priors” workshop paper from MLSB NeurIPS 2025 [43] applies physics-informed priors for general protein–protein interactions but does not address antibody-specific CDR regularization or multi-modal fusion. Loux et al. [44] demonstrated that naïvely adding structural information to ESM-3 can *decrease* prediction accuracy (“More structure, less accuracy”), underscoring that our physics-constrained integration strategy represents a non-obvious solution to multi-modal fusion challenges. Despite these advances over existing methods, several open challenges remain.

### Limitations

Several limitations warrant acknowledgment. First, our GNN embeddings (138 features) exhibited negative transfer under LOCO-DMS evaluation, and the optimized 93-feature model excludes them; more sophisticated SE(3)-equivariant networks may yet improve structural representation [45]. Uncertainty quantification methods beyond ensemble disagreement, such as evidential deep learning or conformal prediction [46, 47], may further improve the confidence protocol. Second, although we integrated ESM-2 embeddings (35M parameters), larger protein language models such as ESM-2 650M [48] or ESM-3 may capture additional evolutionary information. Third, our analysis focused on single-point mutations; combinatorial mutations involving epistatic interactions present additional challenges. Fourth, all evaluated datasets derive from in vitro measurements, which may not fully capture in vivo binding dynamics. Fifth, the cross-pathogen transfer failure we observed (mean 46.7% accuracy, with many transfers below chance level) suggests that pathogen-specific training data remains essential, limiting the applicability of universal binding predictors. Sixth, prospective validation on five held-out complexes achieved mean ensemble MCC of only 0.103, confirming that out-of-distribution complex generalization remains the primary bottleneck [14]. Seventh, Omicron-lineage escape mutations continue to shape the SARS-CoV-2 landscape [37, 49], underscoring the need for routinely updated training data. Finally, all deep learning results require GPU infrastructure (we used NVIDIA L4 via Google Cloud Vertex AI); however, the best-performing classical gradient boosting model runs efficiently on CPU.

## Materials and Methods

### Datasets

**SKEMPI v2.0** [6]: Downloaded from https://life.bsc.es/pid/skempi2/. Binding affinity changes computed as 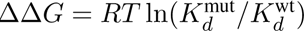 where *RT* = 0.592 kcal/mol at 298K. Binary labels assigned as binding-improving (ΔΔ*G <* 0) or binding-reducing (ΔΔ*G* ≥ 0).

**AbAgym** [11]: Downloaded from https://github.com/3BioCompBio/Abagym. Non-redundant interface mutations (36,541) used, defined as residues within 8 Å of the binding partner. MinMax-normalized DMS scores binarized at the median value.

**CATNAP**: Downloaded from https://www.hiv.lanl.gov/catnap. IC_50_ values log-transformed and binarized at the median for classification.

**AlphaSeq** [3]: Downloaded from https://github.com/mit-ll/AlphaSeq_Antibody_ `Dataset`. A stratified subsample of 50,000 antibodies used for computational tractability.

### Hand-crafted sequence features (111 features)

We extracted 111 hand-crafted features capturing amino acid composition and physicochemical properties:

**Amino acid composition (60 features)**: Fraction of each of the 20 standard amino acids computed separately for VH, VL, and antigen sequences.

**Physicochemical properties (30 features)**: For each sequence (VH, VL, antigen), we computed: mean and standard deviation of hydrophobicity (Kyte-Doolittle scale), charge (+1 for R/K, −1 for D/E), molecular weight, and polarity; net charge; and GRAVY (grand average of hydropathy).

**Sequence length features (5 features)**: VH length, VL length, antigen length, total antibody length, and antigen-to-antibody length ratio.

**CDR region features (14 features)**: For each CDR (VH-CDR1/2/3 and VL-CDR1/2/3), we computed length and mean hydrophobicity. CDR boundaries were defined using approximate positional definitions: VH-CDR1 (residues 26–35), VH-CDR2 (50–65), VH-CDR3 (95–102), VL-CDR1 (24–34), VL-CDR2 (50–56), VL-CDR3 (89–97). Note: these are simplified positional approximations rather than Kabat or IMGT numbering, which require antibody-specific sequence alignment tools.

**Mutation-specific features (2 features)**: Binary indicators for mutation to alanine (alanine scanning) and charge-changing mutations.

### Structural features

PDB structures parsed using BioPython [50]. Interface contacts computed as atom pairs between antibody and antigen chains within 8 Å cutoff using NeighborSearch. Local density computed as the number of contacts divided by 10. Buried classification assigned for residues with *>*5 interface contacts.

### Graph neural network

A three-layer message-passing graph convolutional network (GCN) was implemented operating on the residue contact graph. Nodes represent C*α* atoms; edges connect residues within 8 Å. Node features are 25-dimensional vectors encoding amino acid identity (20-dimensional one-hot), hydrophobicity (Kyte-Doolittle, normalized), formal charge, polarity, aromaticity, and crystallographic B-factor. Message passing with residual connections and LayerNorm updates node features as: 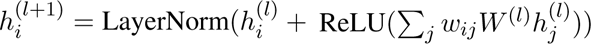 where *w_ij_* are edge weights inversely proportional to C*α* distance and *W* ^(^*^l^*^)^ are learned weight matrices with hidden dimension 64. Dropout (0.3) is applied after each layer. The final representation for each mutation concatenates the mutation-site node embedding (64-dim), a global graph pooling vector (64-dim), and mutation delta features (10-dim) encoding physicochemical property changes, yielding a 138-dimensional embedding per mutation. The network was trained per-LOCO-fold using Adam (lr = 0.001), binary cross-entropy with positive class weighting, and gradient clipping (max norm 1.0) for 30 epochs.

### Classification model

Gradient boosting classifier (scikit-learn) with 100 estimators, maximum depth 5, trained with default hyperparameters. Features standardized to zero mean and unit variance before training.

### Cross-validation and hyperparameter tuning

Three validation strategies were employed:

**Stratified 5-fold CV**: Standard cross-validation preserving class distribution, used for within-dataset evaluation.

**Leave-one-complex-out (LOCO) CV**: All mutations from a single PDB complex held out in each fold (54 folds for SKEMPI), providing stringent evaluation of generalization to unseen protein systems.

**Cross-pathogen evaluation**: Models trained on all mutations from one pathogen and tested on all mutations from a different pathogen.

**Nested CV for hyperparameter tuning**: 5-fold outer loop for evaluation, 3-fold inner loop for hyperparameter selection. Grid search over: Logistic Regression (*C* ∈ {0.01, 0.1, 1}, penalty ∈ {*l*1*, l*2}), Random Forest (n_estimators ∈ {50, 100, 200}, max_depth ∈ {5, 10*, None*}), Gradient Boosting (learning_rate ∈ {0.01, 0.05, 0.1}, max_depth ∈ {3, 5, 7}).

### Evaluation metrics

Classification accuracy, Matthews correlation coefficient (MCC), and area under the receiver operating characteristic curve (AUC-ROC) reported for all experiments. MCC preferred for imbalanced datasets as it accounts for all four confusion matrix categories.

### Code availability

All analysis code is available from the corresponding author upon reasonable request.

### Conclusion

This work demonstrates that multi-scale feature fusion is both necessary and sufficient for antibody-antigen binding affinity prediction to exceed the majority-class baseline under leave-one-complex-out evaluation. The optimized 93-feature representation, combining physicochemical, structural, protein language model, and biophysical descriptors, achieved MCC = 0.206 under LOCO-DMS cross-validation on 36,541 mutations across 68 held-out deep mutational scanning experiments. A confidence-stratified ensemble protocol reached MCC = 0.374 (83.5% accuracy) at 25.5% coverage, approaching 79.1% of the Boltzmann-limited theoretical ceiling imposed by near-neutral mutations. No single feature modality exceeded the 73.3% majority baseline in isolation, establishing that binding affinity prediction is inherently multi-scale. Five deep learning architectures benchmarked under LOCO-DMS for the first time showed self-attention matching classical gradient boosting (both MCC = 0.200), indicating that current feature representations capture the available predictive signal without requiring learned representations. Cross-pathogen transfer failed systematically (mean 46.7%), and prospective validation on five held-out complexes achieved ensemble MCC = 0.103, confirming that generalization to entirely unseen antibody-antigen systems remains the primary open challenge. These results provide calibrated performance expectations for computational antibody engineering and motivate future integration of larger protein language models, SE(3)-equivariant structural representations, and domain adaptation methods for cross-pathogen transfer.

## Acknowledgments

The authors acknowledge computational resources of the Intelligent Robotics and Rebooting Computing Chip Design (INTRINSIC) Laboratory, Centre for SeNSE, Indian Institute of Technology Delhi, IM00002G_RB_SG IoE Fund Grant (NFSG), Indian Institute of Technology Delhi.

## Author contributions

S.S. conceived the study, designed and performed the computational analyses, and wrote the manuscript.

## Data and materials availability

All datasets used in this study are publicly available: SKEMPI v2.0 (https://life.bsc.es/pid/skempi2/), AbAgym (https://github.com/3BioCompBio/ Abagym), CATNAP (https://www.hiv.lanl.gov/catnap), and AlphaSeq (https://github.com/mit-ll/AlphaSeq_Antibody_Dataset). Pre-computed feature matrices and trained model weights are available from the corresponding author upon reasonable request.

